# SARS-CoV-2 variant evolution in the United States: High accumulation of viral mutations over time likely through serial Founder Events and mutational bursts

**DOI:** 10.1101/2021.02.19.431311

**Authors:** Rafail Nikolaos Tasakis, Georgios Samaras, Anna Jamison, Michelle Lee, Alexandra Paulus, Gabrielle Whitehouse, Laurent Verkoczy, F. Nina Papavasiliou, Marilyn Diaz

**Author notes:** These authors contributed equally to this work (Anna Jamison, Michelle Lee, Alexandra Paulus, Gabrielle Whitehouse). Co-corresponding authors, (MD), (LV), (FNP).

## Abstract

Since the first case of COVID-19 in December 2019 in Wuhan, China, SARS-CoV-2 has spread worldwide and within a year has caused 2.29 million deaths globally. With dramatically increasing infection numbers, and the arrival of new variants with increased infectivity, tracking the evolution of its genome is crucial for effectively controlling the pandemic and informing vaccine platform development. Our study explores evolution of SARS-CoV-2 in a representative cohort of sequences covering the entire genome in the United States, through all of 2020 and early 2021. Strikingly, we detected many accumulating Single Nucleotide Variations (SNVs) encoding amino acid changes in the SARS-CoV-2 genome, with a pattern indicative of RNA editing enzymes as major mutators of SARS-CoV-2 genomes. We report three major variants through October of 2020. These revealed 14 key mutations that were found in various combinations among 14 distinct predominant signatures. These signatures likely represent evolutionary lineages of SARS-CoV-2 in the U.S. and reveal clues to its evolution such as a mutational burst in the summer of 2020 likely leading to a homegrown new variant, and a trend towards higher mutational load among viral isolates, but with occasional mutation loss. The last quartile of 2020 revealed a concerning accumulation of mostly novel low frequency replacement mutations in the Spike protein, and a hypermutable glutamine residue near the putative furin cleavage site. Finally, the end of the year data revealed the presence of known variants of concern including B.1.1.7, which has acquired additional Spike mutations. Overall, our results suggest that predominant viral sequences are dynamically evolving over time, with periods of mutational bursts and unabated mutation accumulation. This high level of existing variation, even at low frequencies and especially in the Spike-encoding region may be become problematic when superspreader events, akin to serial Founder Events in evolution, drive these rare mutations to prominence.

**AUTHOR SUMMARY:** The pandemic of coronavirus disease 2019 (COVID-19), caused by the severe acute respiratory syndrome coronavirus 2 (SARS-CoV-2), has caused the death of more than 2.29 million people and continues to be a severe threat internationally. Although simple measures such as social distancing, periodic lockdowns and hygiene protocols were immediately put into force, the infection rates were only temporarily minimized. When infection rates exploded again new variants of the virus began to emerge. Our study focuses on a representative set of sequences from the United States throughout 2020 and early 2021. We show that the driving force behind the variants of public health concern, is widespread infection and superspreader events. In particular, we show accumulation of mutations over time with little loss from genetic drift, including in the Spike region, which could be problematic for vaccines and therapies. This lurking accumulated genetic variation may be a superspreader event from becoming more common and lead to variants that can escape the immune protection provided by the existing vaccines.

## INTRODUCTION

The Severe Acute Respiratory Syndrome Coronavirus 2 (SARS-CoV-2), which causes the Coronavirus disease 2019 (COVID-19), was first detected in December 2019 in Wuhan, China, when a number of severe pneumonia cases were reported [1]. By March 11^th^, 2020, the COVID-19 outbreak was classified as a pandemic by the World Health Organization (WHO) [2] and as of early February 2021, more than 105 million COVID-19 cases have been confirmed worldwide, while 2.29 million related deaths have been reported [3].

SARS-CoV-2 is an enveloped, single-stranded, positive-sense RNA virus and a member of the *betacoronavirus* genera, of the *Coronaviridae* family [4]. The viral envelope of SARS-CoV-2 consists of the membrane (M), envelope (E), nucleocapsid (N) and spike (S) proteins (encoded by the ORF5, ORF4 and ORF2 respectively), crucial components of the viral structure, but also necessary for the packaging of the viral RNA genome, and for viral infectivity [5]. The S protein (also known as Spike glycoprotein), is a major contributor to COVID-19’s pathogenesis and tropism, as it is responsible for SARS-CoV-2’s recognition, fusion and entrance into host cells. The infection process initiates when the Receptor Binding Domain (RBD; S1 subunit) of the S protein recognizes and binds the angiotensin-converting enzyme 2 (ACE2) receptor of the host, leading to fusion of the viral envelope with the cellular membrane thanks to a hydrophobic fusion peptide sequence found in the S2 subunit of Spike [6].

Entrance and subsequent release of the positive strand viral RNA genome in the host cell, is followed directly by its translation into a variety of structural and non-structural proteins crucial for the viral life cycle [7,8]. ORF1a and 1b are the first to be translated and encode the polyproteins pp1a and pp1b, which are cleaved by the papain-like protease (PL^pro^) and the chymotrypsin-like protease (also referred to as 3C-like protease; 3CL^pro^) [9]. This results in the production of 16 non-structural proteins (nsp1-11 from pp1a and nsp12-16 from pp1b) [9]. Together, these nsps are necessary for the viral life cycle as they regulate the assembly or are components of the Replication-Transcription Complex (RTC) [10]. Nsp1 “hijacks” the translational machinery of the host to prioritize viral protein expression [11], while Nsp2 modulates the host’s cell cycle progression, migration, differentiation, apoptosis, and mitochondrial biogenesis [12]. Nsp4 interacts with nsp3 and other host proteins to facilitate viral replication [5,12], while the nsp6 protein induces membrane vesicles [13]. Nsp12 functions as an RNA-directed RNA polymerase (RdRp) and synthesizes the viral RNA with the help of the cofactors nsp7 and nsp8 [14]. Nsp14 is also part of the RTC by virtue of its function as a 3’-5’ exoribonuclease proofreader, among other functions [15]. Additional RTC nsp proteins are nsp9 (capable of binding to RNA), nsp10 (cofactor of nsp14 and nsp16), nsp13 (helicase and 5’ triphosphatase), nsp15 (with N7-methyltransferase function) and nsp16 (with 2’-O-methyltransferase function) [5,16]. Once the RTC complex is established, it produces copies of negative-sense viral RNA, which are then used as templates for synthesis of the positive-sense genomic RNA (through an obligatory double stranded RNA intermediate [17]). These new copies of genomic RNA are either translated for the expression of new nonstructural proteins or are assembled into virions toward viral release [5]. Finally, the N protein binds to the newly synthesized positive-sense genomic RNA in the cytoplasm, forming the ribonucleocapsid, which along with the M, S and E proteins, are transported to the endoplasmic reticulum-Golgi intermediate compartment (ERGIC) for virion assembly. The virions exit the Golgi via budding and are released out of the cell through exocytosis [8].

All these ORFs encode components crucial to the SARS-CoV-2 life cycle. Genomic variants that alter the amino acid composition of any of these ORFs are of interest. Normally such variants would arise from polymerase-induced mutations during viral replication. However, SARS-CoV-2 (with a genome of ~30 kb) appears to mutate less frequently than viruses with smaller genomes [18], a feature attributed to nsp14, which possesses 3’-5’ exoribonuclease proofreading function that repairs some of the RdRp generated errors [15]. Indeed, the majority of single nucleotide variants detected in viral genomes (65% of documented mutations [19,20]) are C-to-U and A-to-G base changes, a likely result of the action of RNA editing deaminases [21]. These enzymes of the APOBEC (Apolipoprotein B mRNA editing enzyme, catalytic polypeptide-like) and ADAR (Adenosine Deaminase Acting on RNA) families are normally referred to as anti-viral [22–24]. They target C’s in single stranded RNA (as is documented for APOBEC1 [25], APOBEC3A [26,27] and possibly APOBEC3G [28]) or A’s in double stranded RNA (generated during viral genome replication–a perfect substrate for ADAR enzymes) to generate transition mutations (C-to-U and A-to-I, decoded as G) [23]. While RNA deamination in general (also referred to as RNA editing) is normally thought of as anti-viral, there is no reason why it cannot power viral evolution as well, and in fact, current data suggest it does so in SARS-CoV-2. Aside from single nucleotide substitutions, there is experimental evidence that, at least *in vitro*, this and earlier coronaviruses (e.g. SARS-1) are capable of recombination, through template strand switching [29].

Here, we have tracked the appearance of mutations in the SARS-CoV-2 genome through the first 12 months of the pandemic in the United States. Starting from aggregate mutational profiles, we derived a number of mutational signatures, representing distinct variants, which we have then tracked as they emerged across the U.S. in the course of the pandemic over time. We report an increase of variant emergence and mutations per variant with time, underscoring the need for continued mitigation even in the context of a successful vaccination strategy. Finally, a few of the variants we identify from early 2021 are evolved versions of the British variant of concern (B1.1.7) further underscoring the urgency of a dual strategy of mitigation and vaccination.

## RESULTS

### SNVs accumulate progressively with time throughout the SARS-CoV-2 genome

The SARS-CoV-2 isolates analyzed in this study come from infected American individuals collected between January 19^th^ 2020 and January 6^th^ 2021 and encompass 8171 sequences. The number of sequences and locations where they are obtained from are shown in Fig S1. Although viral isolate numbers decrease right after the first wave, a second increase in daily isolate numbers starts from late July onwards. The number of SNVs per viral isolate increases progressively over time (Fig 1A), indicating the virus is not keeping a static genomic profile during the course of the pandemic and is instead accumulating diversity.

**Figure 1.**
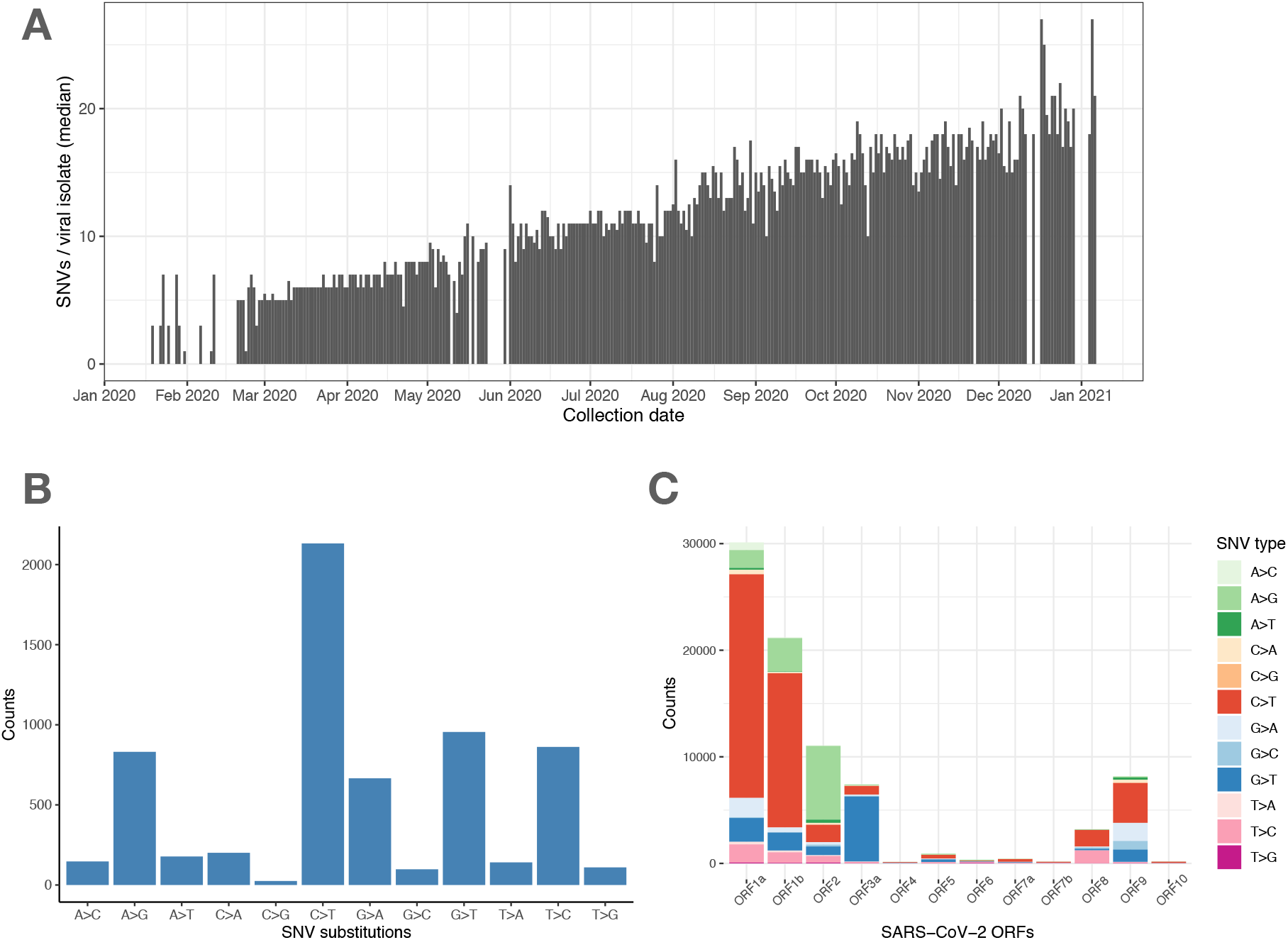
SARS-CoV-2 viral genomes accumulate specific sets of SNVs over time. (**A**) Frequency histogram showing the steady increase of SNVs called per viral isolate over time (Collection Date), indicating their aggregation in SARS-CoV-2 genomes. (**B**) Distribution of substitutions at unique SNVs. Two of the most frequent SNV substitutions, C>T and A>G, have been previously associated with APOBEC and ADAR deaminase activities, on the SARS-CoV-2 ssRNA(+) genome or its dsRNA intermediate, respectively. (**C**) Graphical representation of SNV substitution profiles at various SARS-CoV-2 ORFs, illustrating intrinsic mutational bias for C>T dominating the mutation pattern in some ORF’s (i.e. 1a and 1b), but being masked (likely by selection) in other ORF’s like ORF2 encoding Spike region.

The kinds of substitutions that characterize the aggregate viral SNV profile are predominantly C>T changes, with A>G, G>T and T>C also abundantly represented (Fig 1B) among all mutations. When examining the mutation pattern among unique (non-ancestral) synonymous changes in an effort to better understand the mechanism generating variability in SARS-CoV-2, we found that C to T (U) and T (U) to C transitions are over-represented among synonymous changes (Table 1; synonymous changes typically roughly represent 1/3 of the mutations). The large number of C to U mutations (by far the most common mutation when only unique mutations are considered) regardless of whether they generate a replacement or not, combined with their excess representation among synonymous changes, suggest the intrinsic signature of the mechanism generating mutations in SARS-CoV-2 involves the generation of C to T mutations with a secondary smaller bias for T to C. These base substitution patterns add to the increasing chorus in the literature that the APOBEC family of RNA editing enzymes may be contributing to SARS-CoV-2 diversity (not entirely surprising considering their known roles in antiviral activity [20,21,24]). In certain ORFs, C to T changes were predominant (Fig 1C) while others deviated from the intrinsic mutational signature such as ORF2 encoding spike, suggesting the intrinsic pattern may be masked by positive selection for other mutations in ORF2.

**Table 1.** Nucleotide substitution ratios of synonymous to non-synonymous changes among transtitions, G-to-A, A-to-G, C-to-U, U-to-C.

|  | G-to-A | A-to-G | C-to-U | U-to-C |
| --- | --- | --- | --- | --- |
| Silent (S) | 33 | 52 | 405 | 96 |
| Missense (M) | 81 | 81 | 383 | 32 |
| Ratio (S/M) | 0.41 | 0.64 | 1.06 | 3.0 |

In our viral isolate cohort, fourteen specific missense mutations were found at high frequencies in the aggregate sequence data (Fig 2A; Table 2) suggesting they were under positive selection. Mutations that appear in more than 10% of the retrieved sequences whose frequency over time profile suggest at least three major variants include following:

1. in ORF1a, a Threonine-to-Isoleucine (T>I; T265I) change is present at about 48.79% of the sequences leading to a recoding effect in the Nsp2 protein (also found in [30]), which is one of the first viral encoded proteins to initiate the viral life cycle, as well as a Leucine-to-Phenylalanine (L>F; L3352F) in 12.98% of sequences, recoding the peptidase C30. The frequency of this mutation follows a specific pattern (pattern A. Fig 2B), where it has overall increased over time concurrently with other mutations of the same pattern as mentioned below.
2. In ORF1b, a Proline-to-Leucine (P>L; P4715L in 82.03% of sequences) appears to be the most frequent mutation found in our cohort and has been previously also found in [31]. In the same ORF, Y5865C (Y>C) and P5828L (P>L) represent recoding changes affecting the DNA/RNA helicase domain, and N6054D and R7014C represent amino acid changes in Nsp14 and Nsp16 respectively [32]. Not all of these mutations follow the same frequency patterns (for example P4715L, P5828L and Y5865C, and R7014C follow distinct patterns (Figure 2B)).
3. In ORF2 an Aspartic-acid-to-Glycine (D614G) change (in 80.76% of sequences), (pattern B, Figure 2B), maps between the receptor-binding domain (RBD) and the S2 subunit of the spike. This change has been extensively noted in the literature as a variant associated with an increase in infectivity and appears to have originated in Europe [33].
4. In ORF3a, a Glutamine-to-Histidine (Q>H; Q57H) mutation (~57.62% of the sequences - pattern B, Figure 2B) is also found along with a G172V (G>V) change [34] (pattern A, Figure 2B), both recoding the viroporin protein of SARS-CoV-2 [35].
5. Mutations in the Ig-like (ORF8) and Nucleocapsid (ORF9) proteins of SARS-CoV-2 have also been abundantly found: in the first an S24L, which follows a unique frequency pattern (pattern D, Figure 2B), and an L84S change (pattern A, Figure 2B), both at about 15% frequency) [36] and in the latter a P199L change [37] and a P67S alteration, which has not been previously documented, at about 10% of the sequences (with a frequency pattern A – Figure 2B).

**Figure 2.**
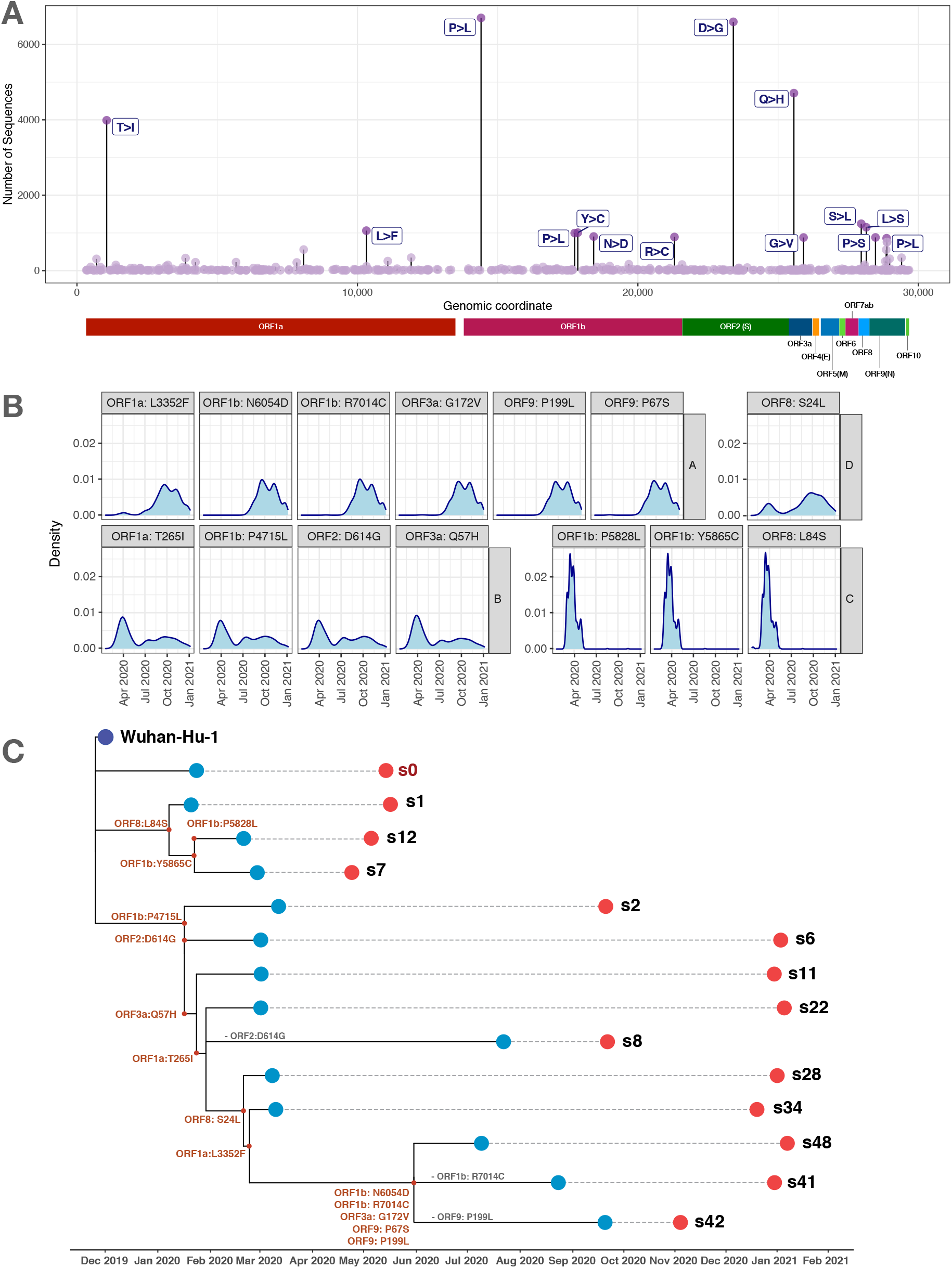
Accumulation of mutations in SARS-CoV-2 genomes and evolution of variants. (**A**) Dot plot representation of missense mutations identified in the SARS2-CoV-2 genome, of which fourteen were found in the most abundant SNVs, including Threonine-to-Isoleucine (T>I) and Proline-to-Leucine (P>L) change in ORF1b (present in about 48.79% and 80.2%) of the sequences, respectively) and the well-documented Aspartic-acid-to-Glycine (D>G) change in ORF2 (found in 80.76% of the sequences). A summary further detailing these predominant mutations is provided in the Table 1 and Supplementary File 1. (**B**) Density histograms (showing how the most common mutations from Fig 2A change over time), reveal that the most common mutations can be grouped into four distinct patterns (A-D); these mutational cooccurrences thus indicate the presence of at least three major variants. (**C**) Unique profiles of co-occurring mutational signatures from the dataset were employed to compile 48 sub-variant putative signatures (s1-s48; Figure S2A) distinct from the original Wuhan viral isolate (s0). 14 signatures and the s0, were found in more than 0.1% of the sequences. Time-scaled phylogenetic tree of those 14 subvariants and s0 (highlighted in red) reveals accumulation of mutations and more complex signatures with an acute burst of mutations in the summer of 2020 likely leading to a novel homegrown variant (s48). The first and last sequences by time profiled (per signature) are denoted with light blue and red dots respectively. The reference genome (Wuhan-Hu-1) is denoted with a dark purple dot. Gain of mutations in the clades is denoted with red letters for each specific mutation, while loss with grey. The most abundant signatures in the end of 2020 and early 2021 are s6, s22 and s48 (also shown in Figure S2B).

**Table 2.**
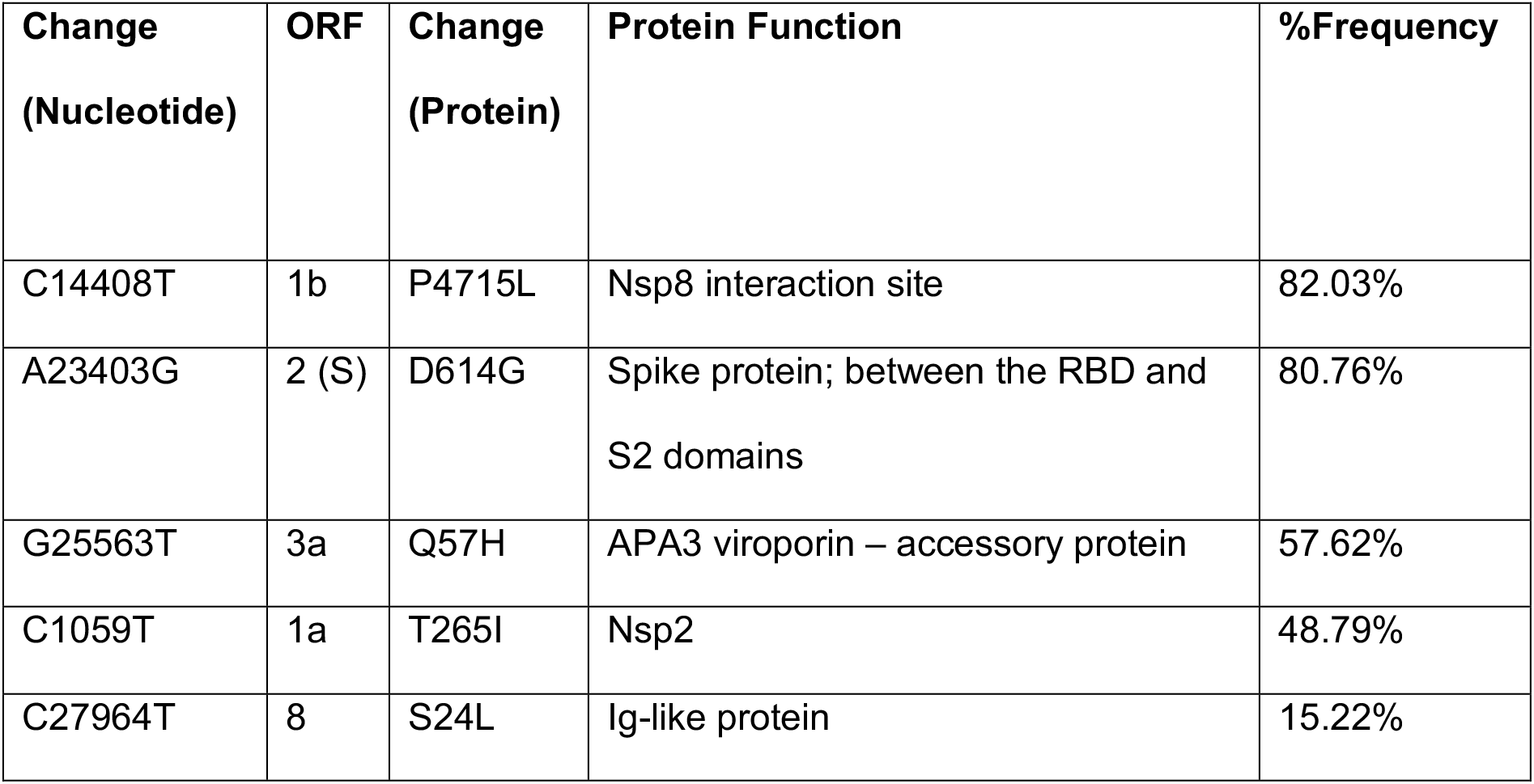
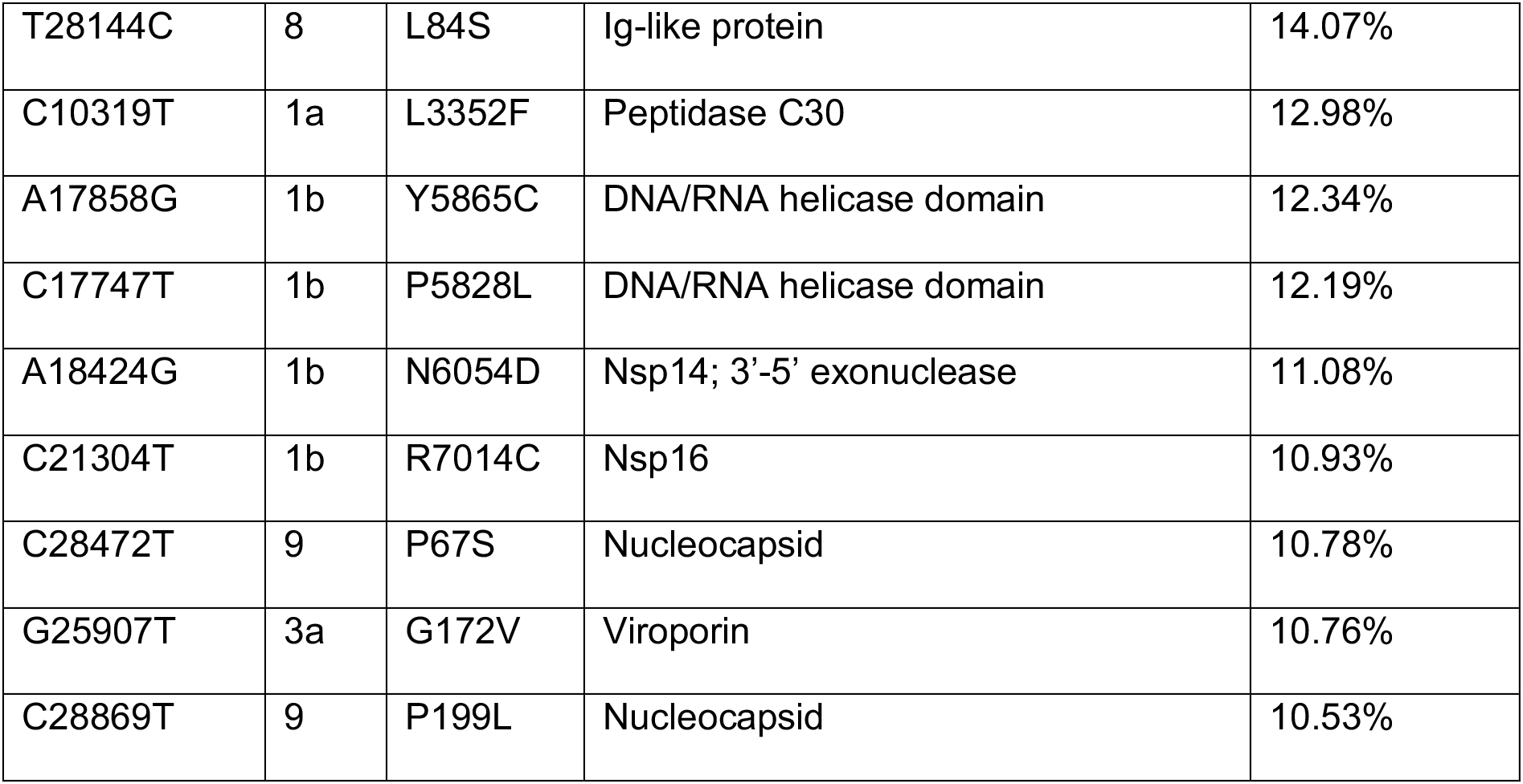
Summary of predominant mutations detected in SARS-CoV-2 genomes. Summary information includes: nucleotide change in position vs. the reference genome, ORF and protein amino acid change, related protein and function that the recoding effect may affect and the percentage frequency (% number of sequences found in). The genomic variants presented in this table are the ones found in more than 10% of the sequences and annotated in figure 2A.

The identified patterns of specific mutation groups with near identical “frequency over time” profiles suggest at least three major variants were present in the United States at various time points in 2020. Some of the mutations are found more frequently earlier in the pandemic rather than later (pattern C; Fig 2B) which correlate with the original Wuhan strain and its early derivatives.

### Mutational signatures over the SARS-CoV-2 genome suggest a combination of genetic drift and selection

From our sequence cohort, we determined all potentially distinct mutation combinations among sequences to get a sense of the evolution of SARS-CoV-2 in the United States. We found 48 distinct putative signatures (s0-s48) that ranged from extremely rare (1 genome) to frequent (in more than 10% of the genomes) (Fig S2A). We focused on those signatures that were present in more than 0.1% of the genomes (Fig 2C). Their prevalence as a function of time was also evaluated (Fig 2C, S2B). Three major variants appear to have dominated the landscape in the US in 2020. These include (a) the reference Wuhan sequence which disappeared as of June 2020, (b) the D to G clade (D614G) and various lower frequency but highly similar subvariants and (c) a group of signatures from that clade that appear to have acquired multiple mutations as a burst event in the summer of 2020 (involving at least 5 missense mutations (Fig 2C)).

These signatures provide enough resolution to examine their distribution across states through time. We focused on states from where sequences were reported both early in the pandemic but also throughout the year. As is clear from Figure 3, significant divergence from the original Wuhan strain is already apparent in mutational profiles of SARS-CoV-2 genomes collected between March and May 2020 (part of the 1^st^ wave). Several mutational signatures become dominant over time and this pattern is specific to some states and seemed to be anchored by the well-known D614G mutation. For example, in California, a diverse set of signatures is present early on, but by the end of 2020, s6, s11, s22, s28 and s48 dominate. Additionally, some signatures are also state-specific such as s41 in MA, and s42 in WI, both very similar to the now ubiquitous s48 but that with the apparent loss of a single mutation in that lineage (Fig 2C), likely through genetic drift. The net effect has been that sequence diversity among viral isolates has increased with time but that diversity may come in bursts as the one seen in the summer of 2020 leading to the s48 signature, likely a homegrown variant (Fig 2C, 3). Intriguingly, one of the mutations that define s48, N6054D, appears to impact the proofreading activity of SARS-CoV-2 [32], raising the possibility that the mutational burst may also be associated with this mutation. These data clearly indicate the genome of SARS-CoV-2 is not static and can adapt through mutation.

**Figure 3.**
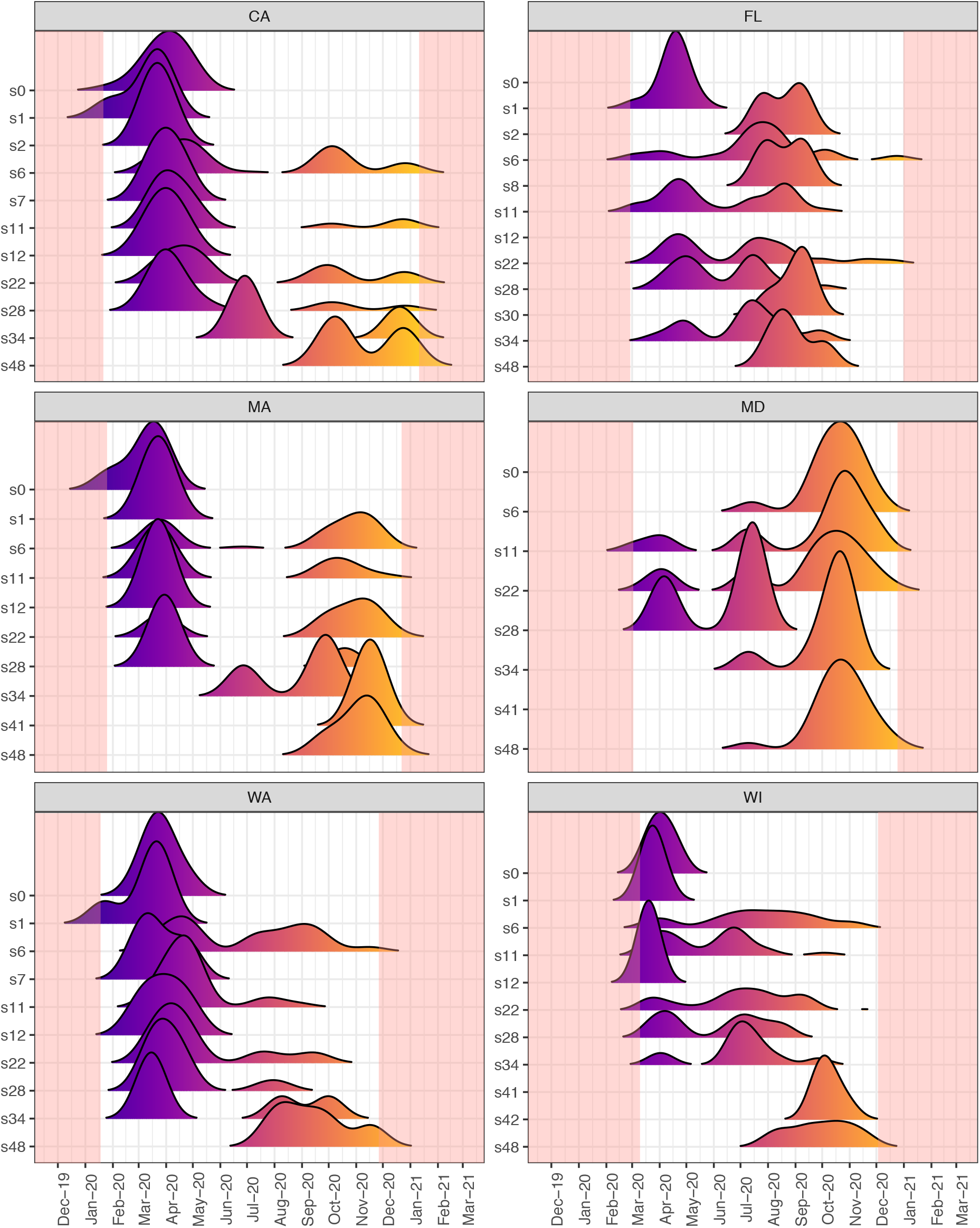
SARS-CoV-2 viral isolate signatures change over time, but with different patterns across states, showing dynamic evolution by mutation, drift and selection. Shown are state-specific ridgeline plots of the density of each signature (y axis) over collection date (x axis). In each plot, peak colors gradually change to highlight transition in time (x axis), with the pink-shaded areas corresponding to the periods of time where data was not available. States shown were selected by sequence abundance throughout the year. Of note, the reference strain s0 is virtually absent by June 2020, while signatures s6, s48, s22 are dominant by late 2020.

### Appearance of SARS-CoV-2 variants of concern in the U.S

The functional consequences of variant evolution are most obvious in the context of Spike protein, as mutations in Spike could impact receptor recognition and infectivity (as well as alter antibody binding and thus lead to immune evasion). Therefore, Spike variants are now denoted as “variants of concern”. One of the first variants of concern was the D614G mutation (clade G) [33], which is now found in the vast majority of SARS-CoV-2 genomes (including all genomes recently annotated as novel variants of concern, such as B.1.1.7). Indeed, D614G is present in more than 80% of the sequences in our cohort in aggregate (Fig2A, Table 2), and virtually all sequences from after the 2^nd^ quartile of 2020 (Q2) have this mutation. In addition to the previously described mutations of concern, we have detected 13 isolates with a H69/V70 deletion. Of these only three have an additional deletion at V143/Y144 (and additional characteristic mutations which define them as B.1.1.7 lineage (Fig 4A-B) according to [38]. The rest carry the H69/V70 deletion together with a handful of other mutations, matching the B.1.375 lineage (Fig 4B). These 13 isolates, detected in the US after mid-November 2020 both in East and West coasts, have continued to evolve. For example, a B.1.1.7 isolate in Florida carries an additional K1191N mutation, and B.1.375 isolates with additional V578L and C1236S mutations have been sequenced from Florida and California respectively. The K1191N mutation in the HR2 domain of B.1.1.7 has been found in at least one other variant in Bangladesh, suggesting this may be another problematic recurrent mutation under positive selection [39]. These findings highlight the ongoing diversification of the Spike region.

**Figure 4.**
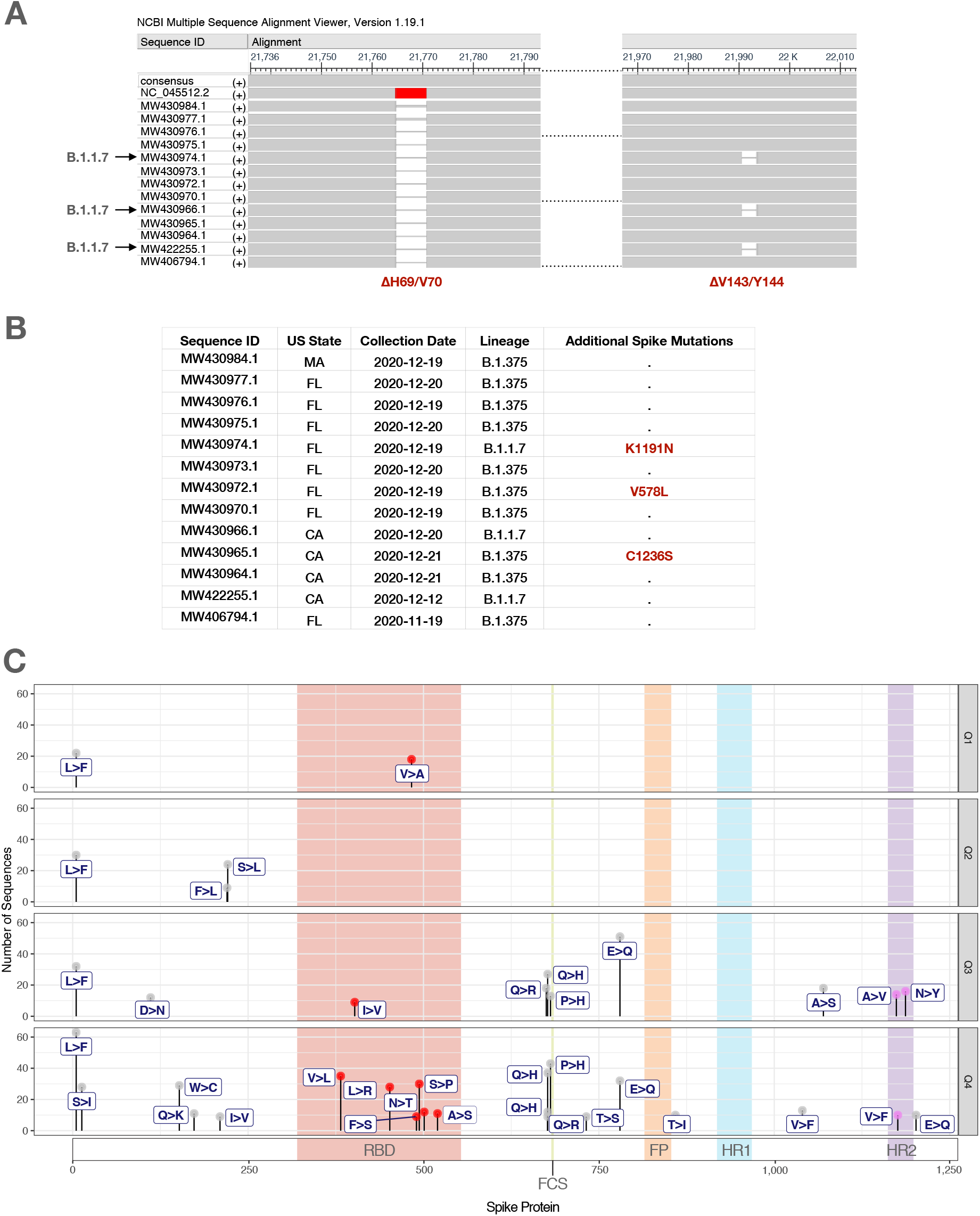
Variants of concern emerging in the United States include novel low frequency mutations in key Spike protein functional domains. (**A**) Sequence Alignment of the Spike protein from 13 isolates in the overall US cohort, carrying the ΔH69/V70 previously described deletion. Of note, three appear to be the B.1.1.7 variant containing the downstream deletion (ΔV143/Y144)(38), first detected in the UK. Also shown are the consensus and reference (NC_045512.2) sequences for the Spike protein. (**B**) Table summarizing further lineage profiling of the 13 isolates shown in Fig 4A, collected from individuals after November 19^th^, 2020 (primarily in CA and FL). All isolates were either B.1.1.7 variants (3), or B.1.375 (10) which lacked the ΔV143/Y144 deletion. Also of note, novel spike mutations were found in a B.1.1.7 lineage isolate in FL (K1191N), as well as in two B.1.375 sequences (V758L and C1236S in FL and CA, respectively). Dot entries in the last column denote the absence of additional mutations. All mutations found in the Spike region of these isolates are summarized in Supplementary Table 1. (**C**) Dot plots showing accumulation of multiple low frequency Spike mutations (LFSM; >0.1% of cohort) in ORF2 over time. The amino acid position per LSFM is shown at the bottom, while quartiles, from the first (Q1) till the last (Q4) are denoted on the right. Mutations in Spike domains are further denoted by shaded areas; RBD/receptor binding domain (red), FCS/furin cleavage site (green), FP/fusion peptide (orange), HR1/heptad repeat region-1 (turquoise) and HR2/heptad repeat region-2 (violet).

### Multiple low frequency missense mutations of unknown consequence are accumulating in the region encoding the Spike protein that may warrant close surveillance

Additional mutations were found in Spike region that are currently at very low frequencies but present in at least 0.1% of sequences (a cutoff selected to minimize sequencing error contribution to the analysis). The consequences of these mutations to infectivity, severity of disease, or response to vaccination remain unknown. They include: L5F (163 genomes), E780Q (83 genomes), P681H (74 genomes) and Q677H (68 genomes; see below), and over 20 additional mostly unidentified amino acid replacing mutations (as summarized in Fig 4C). None of these Spike mutations have been identified as problematic to date, but they remain within the population at very low frequencies. Strikingly, these low frequency spike mutations seem to be increasing over time as more and more mutations accumulate in the 4^th^ quartile - while only a couple have been lost likely from genetic drift (Fig 4C). We found that of these low frequency mutations, six are in the receptor binding domain (RBD) (Fig. 4C) and include (with number of genomes in parentheses): V382L (35), L452R (28), F490S (9), S494P (30), N501T (12), and A520S (11), which may have consequences for binding affinities to the ACE2 receptor in human cells, infectivity and/or response to vaccines developed to trigger antibody responses to the RBD of earlier strains. Moreover, we identified two different amino acid substitutions at Q677: Q677R (A23592G) and Q677H (through two different point mutations −G23593T and G23593C) which are very close to the furin cleavage domain. This hypermutability at Q677 suggests that it is under strong selection. A series of low frequency mostly novel mutations in Spike were detected and included (with number of genomes in parentheses): I210V (9) T719S (9), E781Q (32), T860I (10), V1041F (13), V1176F (10), and E1203Q (10). The entire list of low frequency mutations in Spike are shown in Fig. 4C. Finally, it is important to note that this low frequency variation in Spike currently present within the US population, may be potentially one superspreader event away from prominence, and could lead to problematic new mutations or variants.

## DISCUSSION

Among positive strand RNA viruses, the genome of SARS-CoV-2 has been thought to be remarkably stable - in part because it has proofreading functionality during RNA synthesis - a function carried out by nsp14 [15]. However, this notion of stability has come under scrutiny with the emergence of multiple variants, some threatening the effectiveness of vaccines, and many coinciding in convergently acquired spike mutations [40]. Indeed, though lacking the diversity seen in HIV-1 variants [41], SARS-CoV-2 is fully capable of acquiring mutations that enhance its ability to spread and evade immune responses.

In this study, we aimed to examine SARS-CoV-2 variants in the United States during the first year of the pandemic. We were interested in sampling the existing variation, how variant frequency changes over time and across states and finally, in the potential identification of either new variants or novel mutations in pre-existing variants that have arrived from other parts of the world. We also examined whether the pattern of mutations, particularly among synonymous sites, could provide a clue as to how, despite its proofreading exoribonuclease activity, SARS-CoV-2 has accumulated significant genetic variation.

For this study, we obtained 8171 full length sequences from Covid19 patients from January 2020 through January 2021 from 42 US states. It is important to note that a majority of the data was obtained from a handful of states (California, Florida, New York, Maryland, Massachusetts, Minnesota, Virginia, Wisconsin, and Washington (Fig S1B)), likely a combination of available genome surveillance programs, and rates of infection in those states. We identified several distinct variants (Fig 2C, S2A) that can be categorized as follows: **1)** The original Wuhan strain and a few descendants with minor changes. This strain lacks the D614G change that emerged in Europe early in the pandemic (G-clade). The reference strain and its minor subvariants appear to have been lost in most states by early to mid-summer (Fig. 3); **2)** Two versions of the G-clade European strain defined by the acquisition, in an intermediate within the clade, of multiple mutations within a short period of time in the summer of 2020, leading to the now predominant likely homegrown variant s48 signature (Fig 2C). Our analysis is not compatible with the notion that the burst of mutations originated from a recombination event; rather these mutations appear instead to arise from the acquisition of multiple single base substitutions that increased in frequency in the population relatively quickly, likely through serial Founder Events. However, a few examples of lone mutations shared across variants suggest the possibility of recombination or convergent evolution. Bursts of mutations may also originate from patients with persistent infection despite treatment with convalescent plasma where pressure for immune escape variants may be prolonged and intense [42,43].

Strikingly, the main variants in the US accumulated an increasing number of mutations over time (Fig 1A, 2B-C, 3, S2B). This underscores the fact that with uncontrolled infection, the appearance of new mutations will increase – and this will augment the probability of the emergence of variants that alter the efficacy of vaccines or other therapeutics (e.g. monoclonal antibodies). Of particular concern is our finding that over the last year in the United States, over 20 amino acid replacing mutations arose in the Spike protein that have not been identified yet as problematic, many still remaining in the population but currently at low frequencies (less than 1%, Fig. 4C). Typically, mutations need to reach non-trivial frequency levels to survive genetic drift and loss from the population. However, the number of amino acid replacements impacting ORF2 encoding Spike seems to be increasing over time, with little corresponding loss of variation through drift (Fig. 4C). This low frequency variation in Spike is of concern because of the number of superspreader events in the US population leading to serial Founder Events that can increase the frequency of these rare mutations. Variation reaching non-trivial frequencies through superspreader events can then be subjected to positive selection in viral evolution which can come in the form of host immune escape variants: such as arising in immunocompromised patients receiving convalescent plasma [42,43], or in inadequately vaccinated individuals (e.g. having only received a single dose of a two-dose vaccine regiment), or by outperforming other variants through easier spreading for example. The latter may be the reason the D614G variant became the dominant form in most countries [44,45].

Superspreader events may effectively work as Founder Events in this pandemic. In Founder Events, where a few organisms initiate a new population, typically most genetic variation is lost [46]. However, multiple or serial Founder Events originating from a population can potentiate the generation of new species (or variants in this case) by providing a mechanism for rare mutations to quickly increase their frequency [47]. Therefore, in considering the generation of diversity of SARS-CoV-2, superspreading events is another mechanism, besides mutation, where the virus can effectively increase its diversity over the population. Given this, it is a reasonable possibility that current low frequency Spike mutations may develop into problematic variants through superspreader events. This provides a compelling reason to adhere to strict mitigation controls especially in the context of gatherings with potential to become superspreader events. Minimizing such events is likely critical to control the generation of clinically relevant variants.

Deservedly, a lot of attention has been given to variations in the ORF encoding spike protein (Orf2), since any immune escape mutants are likely to arise particularly (though not exclusively) within the receptor binding domain of Spike, that interacts with the ACE2 receptor [48]. Among variants in Spike, we detected rare instances of the B 1.1.7 variant (3 cases in November in California and Florida) and 10 of the B1.375 variant, recently identified as having the H69/V70 deletion similar to B 1.1.7 but lacking most of the other distinguishing mutations of B.1.1.7 [49]. This novel lineage (B.1.375) is another example of the H69/V70 deletion been found in independent variants, suggesting it evolved convergently multiple times in SARS-CoV-2 variants (as recognized by others [50]) even among different species [51]. Recent models suggest that the H69/V70 deletion may be a gateway alteration to more variation as it may provide increased flexibility of the receptor binding domain to accommodate mutations in the ACE2 receptor among individuals and/or species [48], but this remains speculative. Specific caution is warranted with genomic surveillance of SARS-CoV-2 genomes containing this deletion (whether it is B1.1.7 or not) as it seems to be associated with the generation of new variants of clinical relevance.

In addition to variants within the ORF encoding spike protein (Orf2), we and others found significant variation in other Orfs such as Orf1a and 1b and others, including a 15bp deletion in the region encoding NSP1, previously identified in Japan [52]. The functional relevance of this variation is less clear but cannot be ignored as it may impact the virus ability to replicate, infect other cells once inside the host, and even modulate the host immune response (as has been observed for NSP1 [52]). Therefore, while these non-spike variants may not impact vaccine efficacy, they may impact the severity of the disease and potentially the spread of the virus by increasing its efficiency in hijacking host cells and lowering the viral load threshold required to establish infection.

Because the SARS-CoV-2 genome is not as stable as initially thought despite its proofreading activity, we examined the pattern of mutation among synonymous changes throughout the 8171 SARS-CoV-2 genomes to try to establish the intrinsic signatures of the mechanisms that result in SARS-CoV-2 mutations. Querying synonymous mutations, we found that C-to-U and U-to-C transitions were abundant and among all mutations, C-to-U changes were dominant. One source of C-to-U mutations in RNA is the APOBEC family of RNA editing enzymes, some with anti-viral properties known to deliberately attack the genomes or RNA viruses, such as in the case of Apobec3G and HIV [53]. The less frequent U-to-C mutations could also be due to RNA modification events occurring on uracil, and decoded as cytosine; modifications that could result in such a profile could include thiolation (e.g. 4-thio-uridine) or aminocarboxypropylation (e.g. acp3U) events. Though these modifications have not yet been reported to occur on mRNA, both can occur on tRNA [54, 55]. Finally, A-to-G events are also evident (and the likely result of adenosine deamination to inosine, decoded as guanosine, which is catalyzed by ADAR proteins, whose preference for dsRNA targets could attract them to double-stranded RNA intermediates of the viral replication process) – for example the prominent G clade mutation (D614G) may be the outcome of an A-to-I deamination event at position A23403).

While we are not the first to make the observation that the SARS-CoV-2 genome is a target of modification enzymes [21, 56–58], the fact that (a) many of the mutations that have given rise to variants of concern could be explained by such modification events, together with the fact that (b) modification enzymes have a preference with regard to the nucleotides that neighbor the base-to-be-modified, lead us to speculate that RNA modifications are a major source of targeted mutagenesis of the viral genome. This would explain why emerging variants (like the B1.1.7), rather than diverging in sequence, appear to be acquiring mutations common to unrelated strains (e.g. the new acquisition of the E484K (**G**AA=>**A**AA mutation first defined as concerning in the unrelated 501Y.V2 variant [59], which could be attributed to a modification such as m1G, which can be decoded as A [60]). Overall, this may be good news, as it would imply that the range of sequence alterations that can yield variants of concern is limited, and can be effectively targeted by novel vaccination strategies. This does not eliminate the potential for recombination as another source of variation, as seen often in coronaviruses [29], however we did not detect evidence of recombination events in the sequences we queried here.

A troubling implication of the emergence of Spike variants is the potential for the concurrent development of antibodies that are optimal for the original strains but not for the new strains which may lead to the development of antibody-dependent enhancement (ADE) documented for other viruses and associated with the development of suboptimal antibodies [61]. Fortunately, ADE has not emerged explicitly as a substantial concern with SARS-CoV-2 [62,63], although suspects are the MIS-C or MIS-A, the Kawasaki-like syndromes associated with COVID-19 infection and re-infection both in children and more recently also in adults [64,65].

From a public health perspective, this study underscores the critical importance of mitigating infection levels and particularly, super-spreader events, as critical potential generators of high frequency novel variants from the very low frequency existing and increasing Spike mutation pool. Indeed, the finding of over 20 spike variants at low frequencies in the population of the U.S., is concerning as this “lurking” genetic variation can quickly emerge as novel variants through superspreader, Founder-like events in an expansion process similar to genetic surfing [66,67]. All these considerations require, in addition to recommended mitigation efforts such as social distancing and mask wearing, that large scale vaccination be in accordance to the schedules used in the clinical trials leading to federal agency approval. Further supporting this, are reports of problematic variants arising within individual immunocompromised patients treated with convalescent sera [42] as these escape mutants can clearly arise when subjected to low levels of anti-Spike protein antibody. This, and our finding of potential “lurking” low frequency variants already within the population, dictate that selection against this virus through vaccination be strong and that “taking the foot off the pedal” (for example by allowing people to have a single dose, or by low second shot compliance) can allow this existing variation to give rise to novel escape variants. This explicitly means that vaccinating more people with a single dose, but delaying the second, could lead to suboptimal protection and therefore mild selection pressure on the circulating variants, allowing for the evolution of more robust escape mutants.

## METHODS

### The dataset

The NCBI SARS-CoV-2 Resources portal (https://www.ncbi.nlm.nih.gov/sars-cov-2/) was the source for all SARS-CoV-2 sequences employed in this study. To fulfill the criteria of nucleotide completeness (complete coverage), 8171 viral isolate sequences were retrieved and isolated from human infections in the USA. Sequences retrieved by the time of analysis were isolated from infections reported between January 19^th^, 2020 and January 6^th^, 2021 (noted as “Collection Date” in the database). In our analyses we considered the collection date as the most relevant parameter and interpreted our results according to this time frame.

### Variant calling and annotation

As the reference genome, we considered the one isolated from patient-zero in Wuhan, China (accession number NC_045512 in RefSeq). Alignments were performed with the Multiple Sequence Alignment tool “Clustal Omega” [68,69], comparing each US state separately against the reference genome. We used Clustal alignment outputs (with character counts) as input for our python 3.8 script to call SNVs (Single Nucleotide Variation), which incorporated from ‘biopython’ [70] alignment reading commands for outputs of variation from the reference. Translation of SNVs to note amino acid changes were processed with an R (4.0.2) script, which applied the genetic code on reference sequence to display amino acid variation and thus highlight missense and silent mutations. Annotation of genomic variants with regards to regions in the viral genome (organized into ORFs) was performed employing NCBI RefSeq SARS-CoV-2 genome annotation, which is also publicly available in the NCBI SARS-CoV-2 Resources portal. Most variants and evolutionary signatures called throughout the dataset were visually inspected for validation of SNVs (and presumed amino acid changes). For further analysis and processing different cut-off parameters were followed: as most frequent variants in aggregate (Fig 2A, Table 1) we defined the missense mutations that are present in at least 10% of the sequence cohort (>817 sequences), same cut-off was for the nucleotide changes, but including both silent and missense mutations. For the low frequency Spike mutations, we considered the Spike missense mutations present in more than 0.1% of the sequences.

For detecting specific deletions (eg ΔH69/V70) we employed the BLAST (Basic Local Alignment Search Tool) command line application, by calling for gap-containing sequences compared to the reference, in a locally-constructed database with all viral isolate sequences (n=8171) satisfying the aforementioned criteria.

### Mutational signatures analysis

We defined sequences with distinct combinations of the most frequently detected mutations in SARS-CoV-2 genomes as mutational signatures (Figure 2A; Table 2). All unique combinations were called to build a reference of putative mutational signatures (Fig S2A). We focused on those signatures that were found in more than 0.1% of the sequences (>8). Time-scaled phylogenetic tree of the major signatures (>0.1%) was constructed with IQ-TREE 2 [71].

### Statistical analyses and visualization

All statistical analyses and visualizations were performed in R programming language (v. 4.0.2) employing the R stats package, as well as the Tidyverse (v. 1.3.0) [72], pheatmap (v. 1.0.12), dendextent (v. 1.14.0) [73], msa [74], treeio [75] and ggtree [76].

## AUTHOR CONTRIBUTIONS

This study was conceived by RNT in the summer of 2020 as a summer research project for students from the Nightingale Bamford School, New York. Together with AJ, AP, GW, and ML (a recent alumna), they are responsible for all data collection and analysis. Additional analyses over the fall of 2020 were completed by GS, using the pipelines and scripts established over the summer. MD, LV and FNP provided data interpretation and wrote the manuscript together with GS and RNT.

## ACKNOWLEDGEMENTS

AJ, AP, GW, and ML would like to acknowledge Dr Naomi Kohen of the Nightingale Bamford School and the school’s support for the independent science research program in Heidelberg.

## SUPPORTING INFORMATION

**S1 Figure.** (**A**) Viral sequences isolated from the United States from January 19^th^, 2020 till January 6^th^, 2021 (8171 in total from SARS-CoV-2 NCBI portal; see *Methods*), highest numbers were in mid-to late-March 2020. (**B**) States with the most sequences were WA, WI, VA, CA, MA, FL, MN, MD, NY. Pie chart shows the percentage of sequences per state in from the cohort in aggregate.

**S2 Figure. Associated information with Figures 2 and 3.** (**A**) Heatmap summarizing all the putative signatures (columns) built by the unique combination of dominant mutations (rows). Signatures are ranked from left to right by the sequence number they were found in (red labels below x axis). The first 15 signatures that are found in 0.1% of the sequences (>8) or more were considered for the mutational signature analysis. Presence or absence of mutation in each signature is denoted with blue or light yellow respectively. (**B**) Signatures that were found in more than 0.1% of the viral isolates in aggregate were further explored in the context of time from emergence till early 2021. Viral isolates that were profiled with those signatures (s0, s1-2, s6-8, s11-12, s22, s28, s34, s41-42, s48), were ordered by collection date in the columns of the heatmap from left to right. The heatmap visualizes the % occurrence (light yellow to dark blue scale) of each signature per collection date in the cohort of sequences. Column annotation (bottom of the heatmap) denotes the different quartiles (Q1-Q4) of calendar year 2020 (very few entries from Q1 of 2021 are shown) in which the sequences were collected. The non-variant SARS-CoV-2 (s0) is present primarily in the in Q1 up to mid-Q2 of 2020. While a diverse set of signatures appears in the USA from the start of the pandemic onward, subvariants currently circulating in the American population are variations of s48, s22 and s6 (with some variation per state).

**S1 Table.** Summarized information per viral isolate (rows) detected with deletion ΔH69/V70 alone or in combination with ΔV143/Y144. In the table, for each of the 13 isolates its NCBI accession number as “Sequence ID” is given, along relevant information about the US State or date that the isolate was collected. Dots (“.”) in the last column stand for silent mutations. Mutations marked with bold red are novel ones for their lineage. Associated information with Figures 4B-C.

**S1 File.** Robust list of mutations (silent and missense) detected in the cohort of sequences.

**S2 File.** Low Frequency Spike Mutations (LFSM) detected in the quartiles (Q1-4) of 2020.

