## Supplementary figures and images for "SARS-CoV-2 variant evolution in the United States: High accumulation of viral mutations over time likely through serial Founder Events and mutational bursts"

### S1_Figure

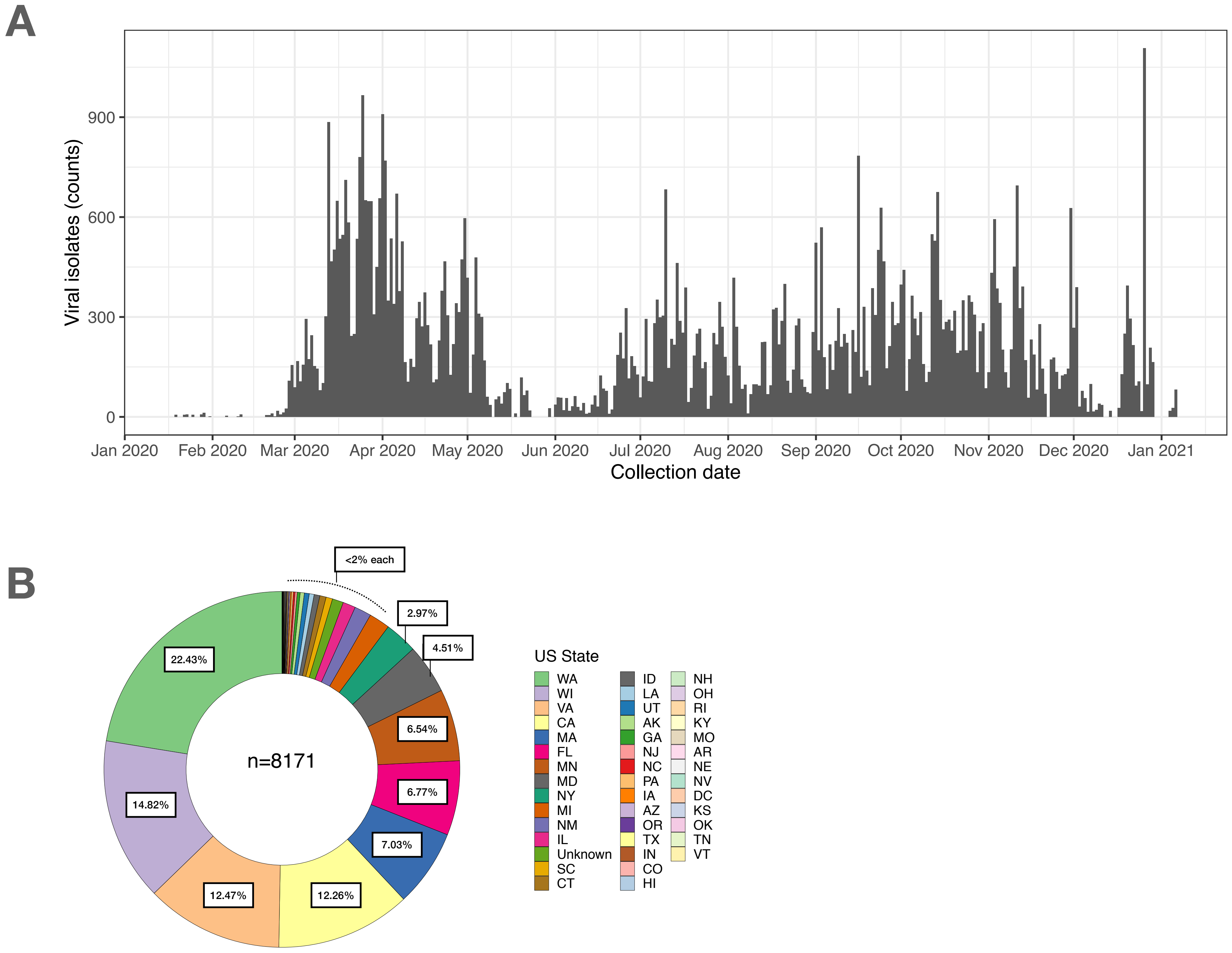

### S2_Figure

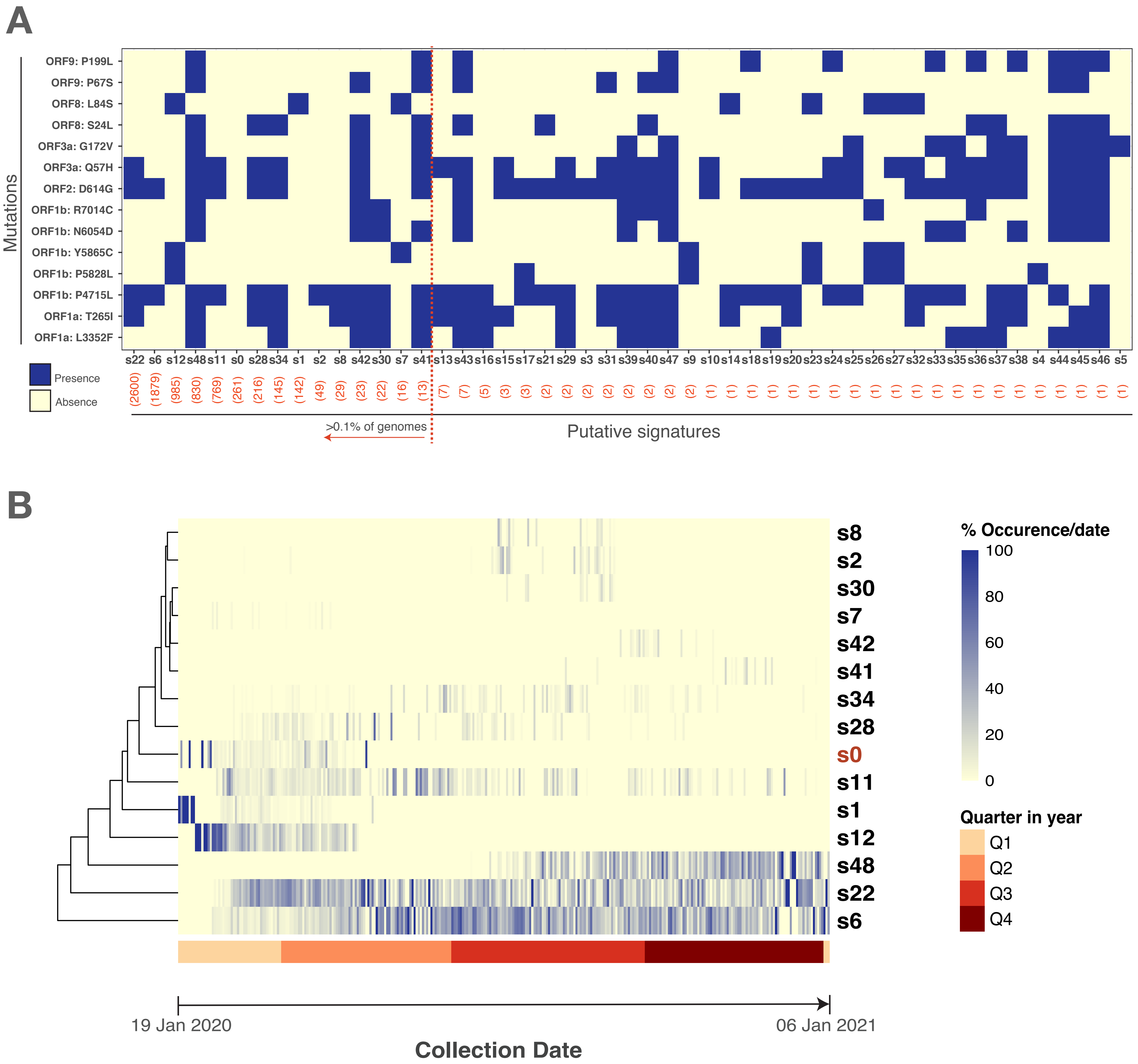
