## Supplementary material for "SARS-CoV-2 variant evolution in the United States: High accumulation of viral mutations over time likely through serial Founder Events and mutational bursts": S1_Table

| **Sequence ID** | **US State** | **Collection Date** | **Lineage** | **Spike Mutations**  **(Nucleotide)** | **Spike Mutations**  **(Protein)** |
| --- | --- | --- | --- | --- | --- |
| MW430984.1 | MA | 2020-12-19 | B.1.375 | C21691T  A23403G  G24007A  C24442T | .  D614G  .  . |
| MW430977.1 | FL | 2020-12-20 | B.1.375 | A23403G  G24007A | D614G  . |
| MW430976.1 | FL | 2020-12-19 | B.1.375 | A23403G  G24007A | D614G  . |
| MW430975.1 | FL | 2020-12-20 | B.1.375 | A23403G  G24007A | D614G  . |
| MW430974.1 | FL | 2020-12-19 | B.1.1.7 | A23063T  C23271A  A23403G  C23604A  C23709T  T24506G  G24914C  **G25135T** | N501Y  A570D  D614G  P681H  T716I  S982A  D1118H  **K1191N** |
| MW430973.1 | FL | 2020-12-20 | B.1.375 | A23403G  G24007A | D614G  . |
| MW430972.1 | FL | 2020-12-19 | B.1.375 | **G22139T**  A23403G  G24007A  C25324T | **V578L**  D614G  .  . |
| MW430970.1 | FL | 2020-12-19 | B.1.375 | C22388T  A23403G  G24007A | .  D614G  . |
| MW430966.1 | CA | 2020-12-20 | B.1.1.7 | A23063T  C23271A  A23403G  C23604A  C23709T  T24506G  G24914C | N501Y  A570D  D614G  P681H  T716I  S982A  D1118H |
| MW430965.1 | CA | 2020-12-21 | B.1.375 | A23403G  G24007A  **T25268A** | D614G  .  **C1236S** |
| MW430964.1 | CA | 2020-12-21 | B.1.375 | T22219C  A23403G  G24007A | .  D614G  . |
| MW422255.1 | CA | 2020-12-12 | B.1.1.7 | A23063T  C23271A  A23403G  C23604A  C23709T  T24506G  G24914C | N501Y  A570D  D614G  P681H  T716I  S982A  D1118H |
| MW406794.1 | FL | 2020-11-19 | B.1.375 | A23403G  G24007A | D614G  . |
